# A versatile fluorescent probe for imaging fibrillar collagen during avian morphogenesis

**DOI:** 10.64898/2026.09.09.750379

**Authors:** Rintaro Tanimoto, Tatsuki Matsui, Junpei Kuroda

## Abstract

Fibrillar collagen architecture provides a fundamental structural framework for vertebrate tissues. Understanding how collagen matrices are organized and remodeled requires high-resolution imaging approaches capable of resolving their three-dimensional architecture and developmental dynamics *in vivo*. We previously established that the fluorescent probe DAF-FM DA enables sensitive visualization of collagen structures in zebrafish, amphibian, and mouse tissues. However, whether this approach is applicable to avian embryos—an important model system in morphogenesis research—has remained unclear. Here we demonstrate that DAF-FM DA specifically and robustly labels collagen matrices in quail embryonic tissues. Using this method, lattice-like stromal networks in the cornea and spongy matrices in the limb cartilage primordium were clearly visualized throughout deep tissue regions. This imaging strategy provides a practical and versatile tool for investigating collagen-matrix development in vertebrate morphogenesis using avian embryos.

## Introduction

Fibrillar collagens, primarily types I and II, are ubiquitous structural proteins in vertebrate tissues (Ricard-Blum, 2011). They are secreted into the extracellular space as triple-helical trimers, where they polymerize into collagen fibrils and assemble into tissue-specific architectures (Mouw et al., 2014; Sherman et al., 2015). These architectures are key determinants of the morphology and function of diverse organs, including skin (Yang et al., 2015), tendon (Franchi et al., 2007), bone (Fratzl et al., 2004; Stockhausen et al., 2021), and cartilage (Bielajew et al., 2020). Thus, the spatiotemporally regulated formation of collagen matrices is indispensable for vertebrate morphogenesis.

Despite this importance, the dynamic processes mediated by cellular interactions—such as collagen production (Mouw et al., 2014; Silverman et al., 2024), fibril growth (Kalson et al., 2015), and matrix degradation (Sprangers and Everts, 2019)—remain poorly understood. Elucidating these mechanisms requires imaging approaches capable of resolving collagen structures *in vivo* with high sensitivity and depth. Despite advances in collagen-fiber imaging techniques, including electron microscopy (Starborg et al., 2008), histochemical methods (Lattouf et al., 2014; Golberg et al., 2024), immunostaining (Yokomizo et al., 2012; Leng et al., 2022; Jacobson et al., 2024), fluorescent protein tagging (Morris et al., 2018; Shiflett et al., 2019; Kashimoto et al., 2022; Hino et al., 2024), and second harmonic generation (SHG) (Han et al., 2005; Aghigh et al., 2023), these approaches are limited in sensitivity, versatility, or technical accessibility. In particular, capturing three-dimensional collagen architecture across whole tissues during development remains challenging.

We recently demonstrated that the fluorescent probe DAF-FM DA (hereafter referred to as DAF in this report) enables highly sensitive visualization of collagen matrices through concise staining procedures in fish (Kuroda et al., 2024; Kuroda et al., 2025; Miyamoto et al., 2026), amphibian (Ohashi et al., 2025; Kuroda et al., 2025), and mammalian tissues (Kuroda et al., 2025).

Because DAF permeates tissues efficiently and covalently reacts with allysine residues in collagen, it uniformly labels fibrillar structures distributed within deep tissue regions (Kuroda et al., 2025). Accordingly, DAF staining achieves faithful three-dimensional collagen imaging in vertebrate tissues, compared with conventional fluorescence-based methods.

Avian embryos, particularly chick and quail, are easy to access and experimentally tractable model organisms (Padgett and Ivey, 1959; Balaban et al., 1988; Hamburger and Hamilton, 1992; Mok et al., 2015). Their developmental and evolutionary significance has also led to their broad use in vertebrate morphogenesis research (Tamura et al., 2011; Egawa et al., 2018; Griffin et al., 2022; Jimenez and Simoes-Costa, 2026). In addition, their large tissue units and well-defined extracellular matrix (ECM) structures, such as skeletal elements (Nakamura et al., 2019b; Bai et al., 2023), provide an excellent system for studying collagen-matrix organization (Roach, 1997; Merrill et al., 2008; Smith et al., 2013; Chen et al., 2015). However, whether DAF-staining is effective in avian embryos has not yet been investigated. Here we show that the DAF-based method robustly resolves the collagen architectures in quail embryonic tissues, notably in the corneal stroma and limb cartilage primordium.

## Materials and methods

### Animal maintenance

Fertilized quail eggs were purchased from Uzuraya (Aichi Prefecture, Japan) and stored at 17°C until use in experiments. For embryo culture, eggs were incubated at 38°C with approximately 80% humidity. All procedures were approved by the Animal Care and Use Committee of Osaka University.

### DAF staining (general procedure)

DAF staining was performed using 5 µM DAF-FM DA (Goryo Chemical, SK1004-01; 5 mM DMSO stock diluted 1:1000 in PBS). Samples were incubated overnight at room temperature unless otherwise noted. After staining, samples were washed with PBS and fixed in 4% PFA overnight at 4°C. Fixed samples were stored in PBS at 4°C until imaging.

### Sample collection

#### Eggshell membrane (ESM)

Eggshells were cracked, and the ESM was peeled off using forceps, cut into small pieces, and washed with PBS. ESM samples were stained following the general DAF procedure. For confocal microscopy, ESM samples were mounted in PBS on an iSpacer (SUNJIN LAB) with either the outer or inner surface facing downward.

#### Embryonic tissues

Embryos at embryonic day 8 (E8) were collected after cracking the eggs. After removing the egg white, yolk, and amnion, embryos were washed three times with PBS and stained following the general DAF procedure.

#### Cornea tissue preparation

Eye tissue was dissected from whole embryos stained with the general DAF procedure, and the cornea was separated from the eyeball using forceps and scalpels. Corneas were subsequently stained overnight at 4°C with phalloidin–Alexa Fluor 594 (Thermo, A12381; 1:100) and Hoechst 33342 (Dojindo, 346-07951; 1:100) in PBS. After washing with PBS, samples were mounted in PBS on an iSpacer with the convex surface facing upward. For optical clearing, corneas were immersed in RapiClear 1.49 (SUNJIN LAB, RC149001) overnight at room temperature and mounted on an iSpacer while immersed in RapiClear.

### Cartilage primordium preparation

#### Ulna

Forelimbs were dissected from whole embryos stained with the general DAF procedure, and surrounding skin and muscle were removed. Ulna samples were separated from forelimbs and subsequently stained overnight at 4°C with phalloidin–Alexa Fluor 594 (Thermo, A12381; 1:100) and Hoechst 33342 (Dojindo, 346-07951; 1:100) in 0.2% PBS-T. Samples were washed with PBS-T and mounted in PBS on an iSpacer.

#### Metacarpal

Using the same procedure as for the ulna, metacarpal samples were subsequently washed with PBS-T, transferred to methanol, and incubated in acetone overnight at -30°C. After washing with PBS-T and returning to PBS, samples were digested with 2 mg/mL hyaluronidase (Nacalai Tesque, 18240-36) for 1 h at 37°C, based on the protocol reported by Francisco Botelho et al., 2015. Samples were blocked in 1 mg/mL BSA in PBS-T for 1 h at room temperature and incubated overnight at 4°C with anti-collagen II antibody (DSHB, II-II6B3; 1:100). The next morning, samples were washed with PBS-T and incubated for 2 h at room temperature with goat anti-mouse IgG–Alexa Fluor 594 (Thermo, A-11032; 1:200). After washing and transferring to PBS, samples were immersed in RapiClear 1.49 overnight and mounted on an iSpacer while immersed in RapiClear.

#### Image acquisition & data analysis

DAF-stained samples mounted on an iSpacer (see sample preparation section) were imaged using an LSM780 confocal microscope (Carl Zeiss) equipped with 10×, 20×, 25× W, and 40× W objectives. Fluorescence signals were detected using the following excitation wavelengths: DAF-FM DA (488 nm), phalloidin (561 nm), and Hoechst (405 nm). For SHG imaging, a two-photon microscope (A1R MP+/Ti2-E, Nikon) equipped with a 20× objective was used with 890 nm excitation and a 440 nm short-pass emission filter. Image visualization and analysis were performed using ZEN 3.5 Blue edition (Carl Zeiss) and Fiji (ImageJ) with the Depth Coding and OrientationJ plugins.

## Results

### Visualization of fibrillar collagen in the eggshell membrane using DAF staining

First, to evaluate the applicability of the DAF-staining method to avian tissues, we examined the eggshell membrane (ESM), a collagen-rich and easily accessible ECM. The ESM is an acellular ECM structure produced by oviduct epithelial cells and consists of two prominent layers —outer and inner—both primarily composed of type I collagen (Wong et al., 1984; Baláž, 2014; Du et al., 2015). Each layer contains a dense, fibrous collagen network (Buss et al., 2023; Pillai et al., 2023). To test whether these fibrous structures can be visualized using the DAF-based method, we peeled ESM samples from quail eggshells, incubated them in DAF solution overnight, and imaged them by confocal microscopy. As shown in Figures 1A and 1B, distinct fibrous collagen structures were clearly visualized in both the outer and inner layers. In the outer layer, fibers up to ∼10 µm in diameter were uniformly distributed, whereas in the inner layer, thinner fibers (∼5 µm) were more densely packed. Orientation analysis further indicated that these fibers lacked a preferred alignment and were arranged randomly in both layers (Figures 1A’ and 1B’). These structural features were consistent with previous descriptions of ESM architecture (Pillai et al., 2023; Buss et al., 2023; Hincke et al., 2000; Preda et al., 2020; Chen et al., 2020). Additionally, the fibrous structures labeled by DAF overlapped with SHG signals, confirming that DAF labels fibrillar collagen in the ESM (Figures 1C–1D’). Together, these results demonstrate that the DAF-staining method reliably visualizes the fibrous collagen structures in the ESM and is applicable to collagen-rich ECMs in avian tissues.

**Figure 1.**
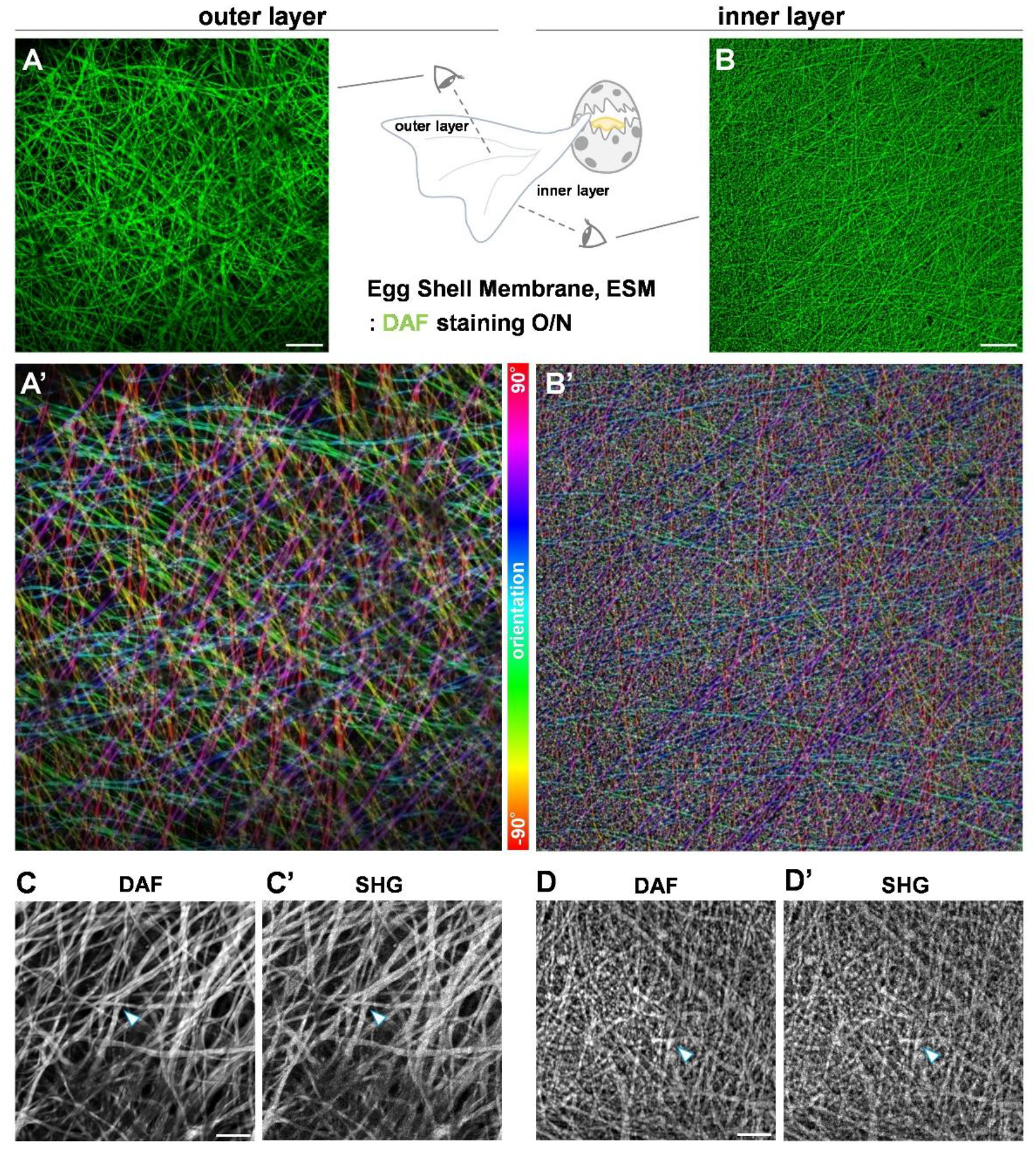
DAF-based imaging of fibrous collagen structures in the eggshell membrane. (A and B) Confocal images of the ESM stained with DAF (green). Scale bars: 50 µm.(A’ and B’) Pseudocolor orientation maps from panels (A) and (B). (C-D’) Multiphoton images of the ESM showing DAF fluorescence and SHG signals. Arrowheads mark regions where DAF and SHG signals well overlap. Scale bars: 20 µm.

### Visualization of stromal fibrillar collagen in the avian cornea using DAF staining

Having confirmed that DAF can stain avian collagen structures, we next examined whether it could label collagen matrices within embryonic tissues. For this purpose, we focused on the cornea, a well-established model for studying collagen-matrix organization (Chen et al., 2015; Lwigale, 2015; Dhouailly et al., 2014). The cornea is particularly suitable because it develops rapidly, is easy to handle, and contains a highly ordered stromal collagen meshwork composed of type I collagen, which underlies its optical and mechanical properties (Meek, 2009; Hassell and Birk, 2010; Quantock et al., 2015).

Quail embryos at embryonic day 8 (E8) were incubated in DAF solution overnight, after which the corneal tissue was dissected and imaged by confocal microscopy. As shown in Figure 2A, stripe-like DAF signals were detected throughout the stromal layer, and color-mapping projection confirmed that this fibrous network extended uniformly across the tissue (Figure 2A’). Higher-magnification imaging revealed fibers 1–3 µm in diameter arranged at regular intervals of 1–10 µm, forming a lattice-like meshwork with minimal fiber crossing (Figure 2C). These features are consistent with the known collagen architecture of the corneal stroma (Meek, 2009; Koudouna et al., 2018; Young et al., 2019). Simultaneously, spindle-shaped keratocytes were interposed between the orthogonally aligned fibers (Figure 2B), and orientation analysis showed that the cellular distribution followed the bimodal orientation pattern of collagen fibers (Figure 2E), in agreement with previous reports (Chen et al., 2015; Koudouna et al., 2018; Young et al., 2014). In addition, the meshwork-like DAF signals overlapped with SHG signals, although SHG signals were also detected in some cells (Figures 2F and 2G), likely reflecting SHG arising from highly oriented actin fibers and other ordered structures (Aghigh et al., 2023). This contrast indicates that DAF preferentially labels collagen fibers within the cornea. These results demonstrated that the DAF-based method specifically resolve the fibrous collagen meshwork in the corneal stroma of avian embryos.

**Figure 2.**
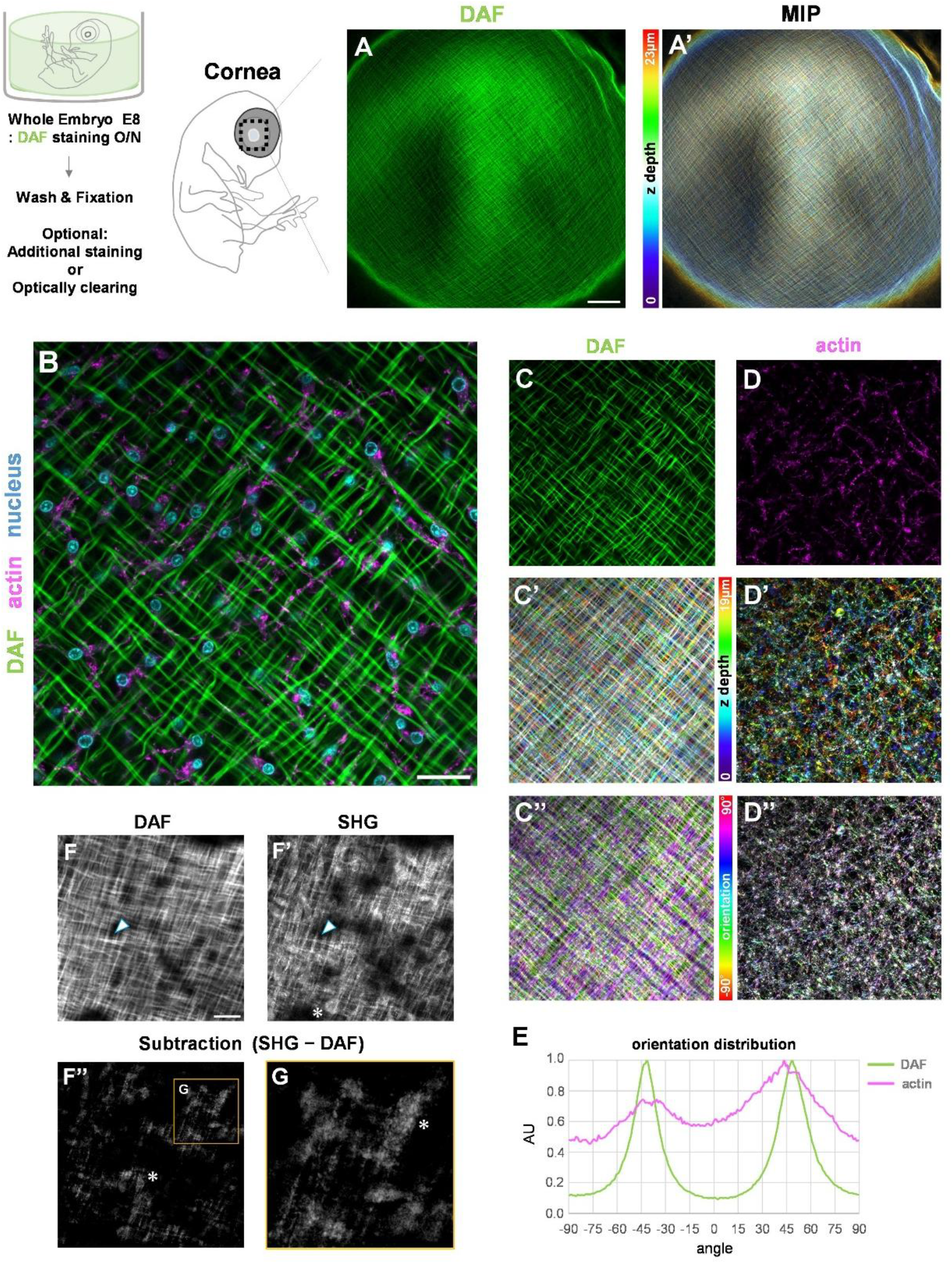
Fibrillar collagen meshwork of the corneal stroma visualized by DAF staining. (A and A’) Confocal images of the cornea tissue stained with DAF (green). (A’) MIP image processed using DepthCoding, 23 µm-stack. MIP: max intensity projection. Scale bar: 100 µm. (B-D) Confocal images of the cornea tissue stained with DAF (green), Phalloidin (magenta) and Hoechst (cyan). Scale bar: 25 µm. (C’ and D’) MIP image processed using DepthCoding, 19 µmstack. (C” and D”) Pseudo-color images processed using OrientationJ from panels (C’) and (D’). (E) Orientation distribution measured from panels (C″) and (D″). (F and F’) Multiphoton images (grayscale) of the cornea tissue stained with DAF. (F’ and G) The subtraction image between panels (F) and (F’). Arrowheads: regions where DAF and SHG signals well overlap. Asterisks: SHG signals from cellular structures. Scale bar: 20 µm.

### DAF-based imaging resolves the porous collagen architecture of the embryonic limb cartilage

Next, we focused on the limb cartilage primordium, which is replaced by bone through endochondral ossification during embryonic development (Nakamura et al., 2019b; Ortega et al., 2004; Yasser et al., 2013; Usami et al., 2016). Limb primordia are suitable for studying collagen dynamics because they extend outward as appendages and are therefore easily accessible for observation (Roach, 1997; Holder, 1978; Tickle, 2004). The cartilage primordium contains abundant in type II collagen and exhibits a characteristic spongy matrix architecture with numerous lacunae (Merrill et al., 2008; Francisco Botelho et al., 2015; Yasser et al., 2013). To evaluate whether DAF can resolve this collagen architecture, we prepared samples using the same protocol used for the cornea, isolated the limb tissue from the trunk, and conducted confocal imaging. As shown in Figure 3A, DAF signals revealed porous structures in the surface region of the ulna primordium. Each lacuna measured tens of micrometers in diameter and contained one or two chondrocytes, consistent with the known organization of spongy cartilage matrices (Holder, 1978; Roach, 1997; Yasser et al., 2013). To assess tissue permeability and sensitivity of the DAF, we removed surrounding tissues from the DAF-stained samples and compared DAF signals with type II collagen immunostaining (Figures 3B–D). In the cross-sectional image of the metacarpal, antibody signals were restricted to approximately 50 µm from the surface, whereas DAF signals were uniformly detected throughout the cartilage and visualized deeper internal structures (Figures 3B’–D’). The fluorescence intensity plot along the cross-section further showed that DAF signals did not exhibit attenuation in deeper regions, indicating uniform labeling without staining unevenness (Figure 3E). The magnified images also demonstrated that DAF signals corresponded well with antibody signals in the surface region, while in deeper regions—where antibody signals were absent—DAF nonetheless revealed clear spongy matrix structures (Figures 3B”–D”). The intensity plot also showed that DAF signals exhibited higher contrast and sharper profiles than antibody staining, suggesting superior detectability of collagen in deeper tissue regions (Figure 3F). Taken together, these results indicate that DAF possesses high tissue permeability and sensitivity in quail embryos, enabling precise visualization of the spatial organization of spongy collagen matrices. Overall, these findings demonstrate that the DAF-based method for collagen-matrix visualization is applicable to avian embryonic tissues in the same way as in other vertebrates (Kuroda et al., 2024; Kuroda et al., 2025; Miyamoto et al., 2026; Ohashi et al., 2025).

**Figure 3.**
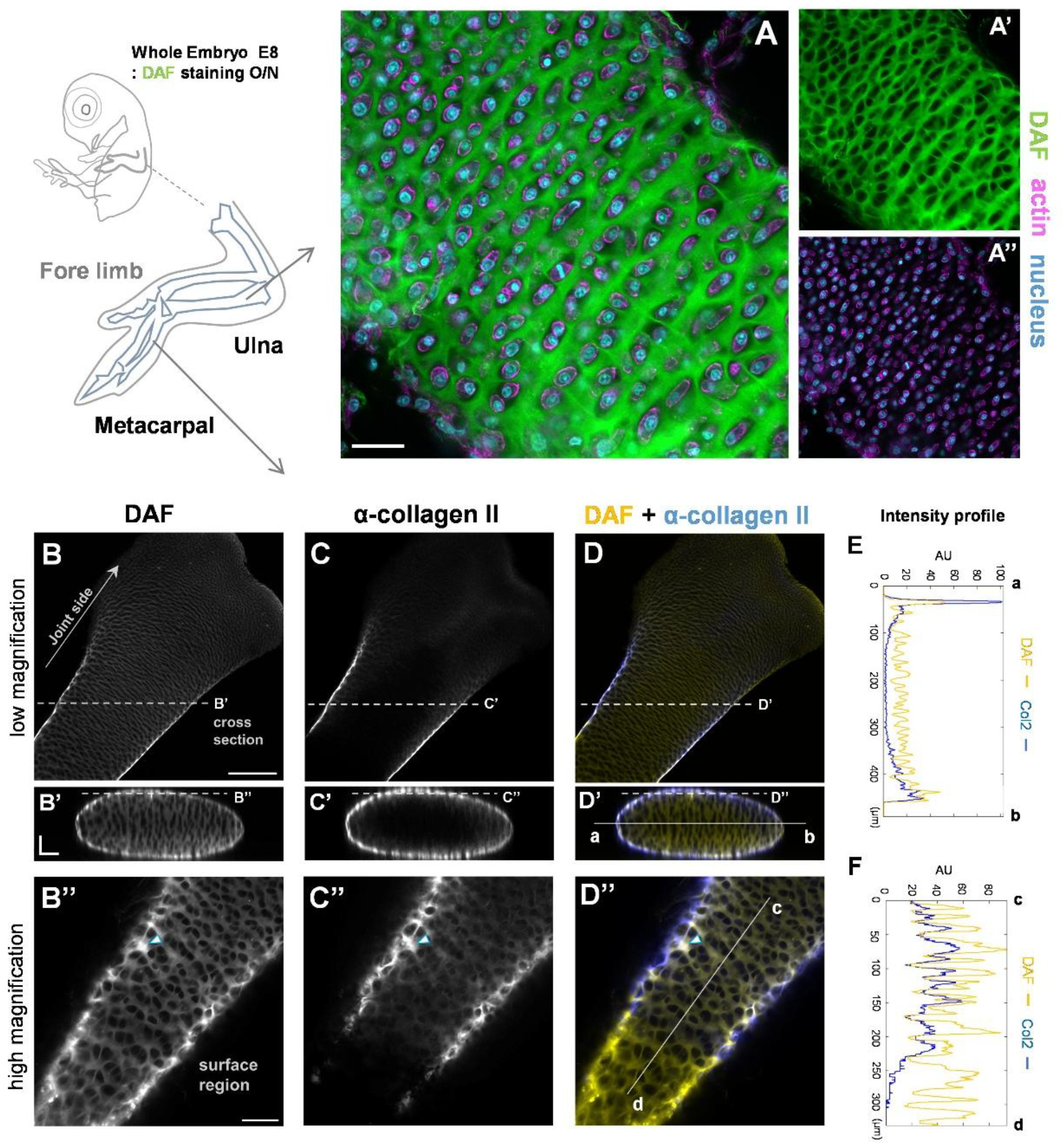
Visualization of the porous collagen matrix in the limb cartilage primordium by DAF staining. (A-A’’) Confocal images of the cartilage tissue stained with DAF (green), Phalloidin (magenta) and Hoechst (cyan). Scale bar: 25 µm. (B-D’’) Confocal images of the cartilage tissue stained with DAF (grayscale or yellow) and anti-collagen II antibody (grayscale or blue). Arrowheads mark regions where DAF and collagen II signals strongly overlap. Scale bars: 150 µm (B), 50 µm (B’ and B”). (E, F) Intensity profiles measured along the dotted lines and solid lines in panels (D’) and (D″).

## Discussion

In this study, we demonstrated that the fluorescent probe DAF-FM DA (DAF) provides highly sensitive labeling of collagen structures in avian embryonic tissues. Notably, the DAF-based method enables clear visualization of three-dimensional collagen architectures, including the lattice-like network in the corneal stroma and the spongy structure in the limb cartilage primordium. These findings provide a foundation for detailed examination of the roles of collagen fibers in corneal stromal development and cartilage-mediated limb skeletogenesis.

Avian embryos develop large tissue units and allow precise staging (Hamburger and Hamilton, 1992; Nakamura et al., 2019a; Bai et al., 2023), providing a powerful model system for analyzing the spatiotemporal assembly of collagen structures (Holder, 1978; Roach, 1997; Young et al., 2014; Hirsinger et al., 2024). However, in vertebrate tissues such as the cornea and cartilage, limited antibody permeability (Yokomizo et al., 2012) and the orientation- and coherency-dependence of SHG (Aghigh et al., 2023) continue to make three-dimensional visualization of collagen matrices challenging. As shown in this study, DAF penetrates whole tissues and yields collagen-specific signals consistent with other markers, thereby complementing existing methods for studying collagen organization.

Although the analyses in this study were performed on fixed samples, the DAF-based method is applicable to living tissues in other vertebrates (Kuroda et al., 2024; Ohashi et al., 2025). In the future, combining DAF staining with in ovo injection (Roto et al., 2016; Cooper et al., 2023) or ex ovo culture systems (Smith et al., 2013; Sukparangsi et al., 2022), will enable tracking of dynamic processes such as collagen-matrix formation, movement, and degradation. Specifically, establishing these techniques is expected to provide novel insights into how collagen matrices are configured, including the lattice-patterned network in the corneal stroma (Chen et al., 2015; Quantock et al., 2015; Young et al., 2019; Koudouna et al., 2023) and the spongy structure in the cartilage primordium (Holder, 1978; Roach, 1997; Smith et al., 2013; Yasser et al., 2013).

Together, the DAF-staining method is effective for avian embryonic tissues and open new avenues for unveiling vertebrate ECM formation mechanisms. As its applications continue to expand, this approach will contribute to understanding a wide range of biological phenomena involving the collagen matrix, including development, wound healing, regeneration and disease.

## Statements

### Data availability statement

The raw imaging data supporting the conclusions of this article are available from the corresponding author upon reasonable request. No custom code was used in this study.

### Ethics statement

All animal procedures were conducted in accordance with institutional guidelines and were approved by the Animal Care and Use Committee of Osaka University. Fertilized quail eggs were obtained and handled following standard ethical regulations for avian embryonic research.

### Author contributions

RT: conceptualization, data curation, formal analysis, investigation, methodology, resources, visualization, writing–original draft, and writing–review and editing. TM: conceptualization, data curation, formal analysis, investigation, methodology, resources. JK: conceptualization, data curation, investigation, methodology, resources, writing–original draft, and writing–review and editing.

## Acknowledgments

We thank the members of the Kondo Laboratory, Graduate School of Frontier Biosciences, Osaka University. We also thank Dr. Ritsuko Suyama, Dr. Yasuhiro Hirano, Dr. Ryosuke Kaneko, and Dr. Ritsuko Morita for their assistance with confocal microscopes at the Graduate School of Frontier Biosciences, Osaka University.

## References

Aghigh, A., Bancelin, S., Rivard, M., Pinsard, M., Ibrahim, H., and Légaré, F. (2023). Second harmonic generation microscopy: a powerful tool for bio-imaging. Biophys. Rev. 15, 43–70. doi: 10.1007/s12551-022-01041-6

Bai, S., Li, S., Li, X., Zhu, S., Shan, Z., Zhang, J., et al. (2023). Comparison of embryonic development, from HH21 to HH40, between ostrich (Struthio camelus) and chicken (Gallus gallus). Dev. Dyn. 252, 668–681. doi: 10.1002/dvdy.568

Balaban, E., Teillet, M.-A., and Le Douarin, N. (1988). Application of the Quail-Chick Chimera System to the Study of Brain Development and Behavior. Science 241, 1339–1342. doi: 10.1126/science.3413496

Baláž, M. (2014). Eggshell membrane biomaterial as a platform for applications in materials science. Acta Biomater. 10, 3827–3843. doi: 10.1016/j.actbio.2014.03.020

Bielajew, B. J., Hu, J. C., and Athanasiou, K. A. (2020). Collagen: quantification, biomechanics and role of minor subtypes in cartilage. Nat. Rev. Mater. 5, 730–747. doi: 10.1038/s41578-020-0213-1

Buss, D. J., Reznikov, N., and McKee, M. D. (2023). Attaching organic fibers to mineral: The case of the avian eggshell. iScience 26, 108425. doi: 10.1016/j.isci.2023.108425

Chen, S., Mienaltowski, M. J., and Birk, D. E. (2015). Regulation of corneal stroma extracellular matrix assembly. Exp. Eye Res. 133, 69–80. doi: 10.1016/j.exer.2014.08.001

Chen, S., Yang, C., Wu, L., Su, H., Zhu, Y., Li, G., et al. (2020). Bioinspired multilevel interconnected networks with porous multiwalled nanotubes built by heterogeneous nanocrystallites. J. Am. Ceram. Soc. 103, 604–613. doi: 10.1111/jace.16698

Cooper, R. L., Santos-Durán, G., and Milinkovitch, M. C. (2023). Protocol for the rapid intravenous in ovo injection of developing amniote embryos. STAR Protoc. 4, 102324. doi: 10.1016/j.xpro.2023.102324

Dhouailly, D., Pearton, D. J., and Michon, F. (2014). The vertebrate corneal epithelium: From early specification to constant renewal. Dev. Dyn. 243, 1226–1241. doi: 10.1002/dvdy.24179

Du, J., Hincke, M. T., Rose-Martel, M., Hennequet-Antier, C., Brionne, A., Cogburn, L. A., et al. (2015). Identifying specific proteins involved in eggshell membrane formation using gene expression analysis and bioinformatics. BMC Genomics 16, 792. doi: 10.1186/s12864-015-2013-3

Egawa, S., Saito, D., Abe, G., Tamura, K. (2018) Morphogenetic mechanism of the acquisition of the dinosaur-type acetabulum. R. Soc. Open Sci. 5 (10), 180604. 10.1098/rsos.180604

Franchi, M., Trirè, A., Quaranta, M., Orsini, E., and Ottani, V. (2007). Collagen Structure of Tendon Relates to Function. Sci. World J. 7, 404–420. doi: 10.1100/tsw.2007.92

Francisco Botelho, J., Smith-Paredes, D., Soto-Acuña, S., Mpodozis, J., Palma, V., and Vargas, A. O. (2015). Skeletal plasticity in response to embryonic muscular activity underlies the development and evolution of the perching digit of birds. Sci. Rep. 5, 9840. doi: 10.1038/srep09840

Fratzl, P., Gupta, H. S., Paschalis, E. P., and Roschger, P. (2004). Structure and mechanical quality of the collagen–mineral nano-composite in bone. J. Mater. Chem. 14, 2115–2123. doi: 10.1039/B402005G

Griffin, C.T., Botelho, J.F., Hanson, M., Fabbri, M., Smith-Paredes, D., Carney, M.R., et al. (2022). The developing bird pelvis passes through ancestral dinosaurian conditions. Nature 608, 346–352. doi: 10.1038/s41586-022-04982-w

Golberg, M., Kobos, J., Clarke, E., Bajaka, A., Smędra, A., Balawender, K., et al. (2024). Application of histochemical stains in anatomical research: A brief overview of the methods. Transl. Res. Anat. 35, 100294. doi: 10.1016/j.tria.2024.100294

Hamburger, V., and Hamilton, H. L. (1992). A series of normal stages in the development of the chick embryo. Dev. Dyn. 195, 231–272. doi: 10.1002/aja.1001950404

Han, M., Giese, G., and Bille, J. F. (2005). Second harmonic generation imaging of collagen fibrils in cornea and sclera. Opt. Express 13, 5791. doi: 10.1364/OPEX.13.005791

Hassell, J. R., and Birk, D. E. (2010). The Molecular Basis of Corneal Transparency. Exp. Eye Res. 91, 326–335. doi: 10.1016/j.exer.2010.06.021

Hincke, M. T., Gautron, J., Panheleux, M., Garcia-Ruiz, J., McKee, M. D., and Nys, Y. (2000). Identification and localization of lysozyme as a component of eggshell membranes and eggshell matrix. Matrix Biol. 19, 443–453. doi: 10.1016/S0945-053X(00)00095-0

Hino, H., Kondo, S., and Kuroda, J. (2024). In vivo imaging of bone collagen dynamics in zebrafish. Bone Rep. 20, 101748. doi: 10.1016/j.bonr.2024.101748

Hirsinger, E., Blavet, C., Bonnin, M.-A., Bellenger, L., Gharsalli, T., and Duprez, D. (2024). Limb connective tissue is organized in a continuum of promiscuous fibroblast identities during development. iScience 27, 110305. doi: 10.1016/j.isci.2024.110305

Holder, N. (1978). The onset of osteogenesis in the developing chick limb. Development 44, 15–29. doi: 10.1242/dev.44.1.15

Jacobson, K. R., Saleh, A. M., Lipp, S. N., Tian, C., Watson, A. R., Luetkemeyer, C. M., et al. (2024). Extracellular matrix protein composition dynamically changes during murine forelimb development. iScience 27, 108838. doi: 10.1016/j.isci.2024.108838

Jimenez, N. A., and Simoes-Costa, M. (2026). The chicken embryo as a model for developmental genomics. Dev. Biol., S0012160626000977. doi: 10.1016/j.ydbio.2026.04.016

Kalson, N. S., Lu, Y., Taylor, S. H., Starborg, T., Holmes, D. F., and Kadler, K. E. (2015). A structure-based extracellular matrix expansion mechanism of fibrous tissue growth. eLife 4, e05958. doi: 10.7554/eLife.05958

Kashimoto, R., Furukawa, S., Yamamoto, S., Kamei, Y., Sakamoto, J., Nonaka, S., et al. (2022). Lattice-patterned collagen fibers and their dynamics in axolotl skin regeneration. iScience 25, 104524. doi: 10.1016/j.isci.2022.104524

Koudouna, E., Mikula, E., Brown, D. J., Young, R. D., Quantock, A. J., and Jester, J. V. (2018). Cell regulation of collagen fibril macrostructure during corneal morphogenesis. Acta Biomater. 79, 96–112. doi: 10.1016/j.actbio.2018.08.017

Koudouna, E., Young, R. D., Quantock, A. J., and Ralphs, J. R. (2023). Developmental Changes in Patterns of Distribution of Fibronectin and Tenascin-C in the Chicken Cornea: Evidence for Distinct and Independent Functions during Corneal Development and Morphogenesis. Int. J. Mol. Sci. 24, 3555. doi: 10.3390/ijms24043555

Kuroda, J., Fujii, K. K., Futaki, S., Hirata, A., Taga, Y., and Koide, T. (2025). A facile method for fluorescent visualization of newly synthesized fibrous collagen by capturing allysine aldehyde groups as cross-link precursors. bioRxiv. 2025.06.19.660320. doi: 10.1101/2025.06.19.660320.

Kuroda, J., Hino, H., and Kondo, S. (2024). Dynamics of actinotrichia, fibrous collagen structures in zebrafish fin tissues, unveiled by novel fluorescent probes. PNAS Nexus 3, pgae266. doi: 10.1093/pnasnexus/pgae266

Lattouf, R., Younes, R., Lutomski, D., Naaman, N., Godeau, G., Senni, K., et al. (2014). Picrosirius Red Staining: A Useful Tool to Appraise Collagen Networks in Normal and Pathological Tissues. J. Histochem. Cytochem. 62, 751–758. doi: 10.1369/0022155414545787

Leng, Y., Lipp, S. N., Bu, Y., Larson, H., Jacobson, K. R., and Calve, S. (2025). Extracellular matrix deposition precedes muscle-tendon integration during murine forelimb morphogenesis. Commun. Biol. 8, 1202. doi: 10.1101/2022.01.23.477427

Lwigale, P. Y. (2015). “Corneal Development,” in Progress in Molecular Biology and Translational Science, (Elsevier), 43–59. doi: 10.1016/bs.pmbts.2015.04.003

Meek, K. M. (2009). Corneal collagen—its role in maintaining corneal shape and transparency. Biophys. Rev. 1, 83–93. doi: 10.1007/s12551-009-0011-x

Merrill, A. E., Eames, B. F., Weston, S. J., Heath, T., and Schneider, R. A. (2008). Mesenchyme-dependent BMP signaling directs the timing of mandibular osteogenesis. Development 135, 1223–1234. doi: 10.1242/dev.015933

Miyamoto, K., Kuroda, J., Kamimura, S., Sasano, Y., Abe, G., Ansai, S., et al. (2026). Actinotrichia-independent developmental mechanisms of spiny rays facilitate the morphological diversification of Acanthomorpha fish fins. Nat. Commun. 17, 2775. doi: 10.1038/s41467-026-69180-y.

Mok, G. F., Alrefaei, A. F., McColl, J., Grocott, T., and Münsterberg, A. (2015). “Chicken as a Developmental Model,” in Encyclopedia of Life Sciences, (Wiley), 1–8. doi: 10.1002/9780470015902.a0021543

Morris, J. L., Cross, S. J., Lu, Y., Kadler, K. E., Lu, Y., Dallas, S. L., et al. (2018). Live imaging of collagen deposition during skin development and repair in a collagen I – GFP fusion transgenic zebrafish line. Dev. Biol. 441, 4–11. doi: 10.1016/j.ydbio.2018.06.001

Mouw, J. K., Ou, G., and Weaver, V. M. (2014). Extracellular matrix assembly: a multiscale deconstruction. Nat. Rev. Mol. Cell Biol. 15, 771–785. doi: 10.1038/nrm3902

Nakamura, Y., Nakane, Y., and Tsudzuki, M. (2019a). Developmental stages of the blue-breasted quail (Coturnix chinensis). Anim. Sci. J. 90, 35–48. doi: 10.1111/asj.13119

Nakamura, Y., Nakane, Y., and Tsudzuki, M. (2019b). Skeletal development in blue-breasted quail embryos. Anim. Sci. J. 90, 353–365. doi: 10.1111/asj.13159

Ohashi, A., Sakamoto, H., Kuroda, J., Kondo, Y., Kamei, Y., Nonaka, S., et al. (2025). Keratinocyte-driven dermal collagen formation in the axolotl skin. Nat. Commun. 16, 1757. doi: 10.1038/s41467-025-57055-7

Ortega, N., Behonick, D. J., and Werb, Z. (2004). Matrix remodeling during endochondral ossification. Trends Cell Biol. 14, 86–93. doi: 10.1016/j.tcb.2003.12.003

Padgett, C. A., and Ivey, W. D. (1959). Coturnix Quail as a Laboratory Research Animal. Science 129, 267–268. doi: 10.1126/science.129.3344.267

Pillai, M. M., Saha, R., and Tayalia, P. (2023). Avian eggshell membrane as a material for tissue engineering: A review. J. Mater. Sci. 58, 6865–6886. doi: 10.1007/s10853-023-08434-2

Preda, N., Costas, A., Beregoi, M., Apostol, N., Kuncser, A., Curutiu, C., et al. (2020). Functionalization of eggshell membranes with CuO–ZnO based p–n junctions for visible light induced antibacterial activity against Escherichia coli. Sci. Rep. 10, 20960. doi: 10.1038/s41598-020-78005-x

Quantock, A. J., Winkler, M., Parfitt, G. J., Young, R. D., Brown, D. J., Boote, C., et al. (2015). From nano to macro: Studying the hierarchical structure of the corneal extracellular matrix. Exp. Eye Res. 133, 81–99. doi: 10.1016/j.exer.2014.07.018

Ricard-Blum, S. (2011). The Collagen Family. Cold Spring Harb. Perspect. Biol. 3, a004978. doi: 10.1101/cshperspect.a004978

Roach, H. I. (1997). New Aspects of Endochondral Ossification in the Chick: Chondrocyte Apoptosis, Bone Formation by Former Chondrocytes, and Acid Phosphatase Activity in the Endochondral Bone Matrix. J. Bone Miner. Res. 12, 795–805. doi: 10.1359/jbmr.1997.12.5.795

Roto, S. M., Kwon, Y. M., and Ricke, S. C. (2016). Applications of In Ovo Technique for the Optimal Development of the Gastrointestinal Tract and the Potential Influence on the Establishment of Its Microbiome in Poultry. Front. Vet. Sci. 3. doi: 10.3389/fvets.2016.00063

Sherman, V. R., Yang, W., and Meyers, M. A. (2015). The materials science of collagen. J. Mech. Behav. Biomed. Mater. 52, 22–50. doi: 10.1016/j.jmbbm.2015.05.023

Shiflett, L. A., Tiede-Lewis, L. M., Xie, Y., Lu, Y., Ray, E. C., and Dallas, S. L. (2019). Collagen Dynamics During the Process of Osteocyte Embedding and Mineralization. Front. Cell Dev. Biol. 7, 178. doi: 10.3389/fcell.2019.00178

Silverman, A. A., Olszewski, J. D., Siadat, S. M., and Ruberti, J. W. (2024). Tension in the ranks: Cooperative cell contractions drive force-dependent collagen assembly in human fibroblast culture. Matter 7, 1533–1557. doi: 10.1016/j.matt.2024.01.023

Smith EL, Kanczler JM, Oreffo RO. A (2013). new take on an old story: chick limb organ culture for skeletal niche development and regenerative medicine evaluation. Eur Cell Mater. 26, 91–106. doi: 10.22203/ecm.v026a07.

Sprangers, S., and Everts, V. (2019). Molecular pathways of cell-mediated degradation of fibrillar collagen. Matrix Biol. 75–76, 190–200. doi: 10.1016/j.matbio.2017.11.008

Starborg, T., Lu, Y., Kadler, K. E., and Holmes, D. F. (2008). “Chapter 17 Electron Microscopy of Collagen Fibril Structure In Vitro and In Vivo Including Three-Dimensional Reconstruction,” in Methods in Cell Biology, (Elsevier), 319–345. doi: 10.1016/S0091-679X(08)00417-2

Stockhausen, K. E., Qwamizadeh, M., Wölfel, E. M., Hemmatian, H., Fiedler, I. A. K., Flenner, S., et al. (2021). Collagen Fiber Orientation Is Coupled with Specific Nano-Compositional Patterns in Dark and Bright Osteons Modulating Their Biomechanical Properties. ACS Nano 15, 455–467. doi: 10.1021/acsnano.0c04786

Sukparangsi, W., Thongphakdee, A., and Intarapat, S. (2022). Avian Embryonic Culture: A Perspective of In Ovo to Ex Ovo and In Vitro Studies. Front. Physiol. 13, 903491. doi: 10.3389/fphys.2022.903491

Tamura, K., Nomura, N., Seki, R., Yonei-Tamura, S., Yokoyama, H., et al. (2011). Embryological Evidence Identifies Wing Digits in Birds as Digits 1, 2, and 3. Science 331, 753–757. doi: 10.1126/science.1198229

Tickle, C. (2004). The contribution of chicken embryology to the understanding of vertebrate limb development. Mech. Dev. 121, 1019–1029. doi: 10.1016/j.mod.2004.05.015

Usami, Y., Gunawardena, A. T., Iwamoto, M., and Enomoto-Iwamoto, M. (2016). Wnt signaling in cartilage development and diseases: lessons from animal studies. Lab. Invest. 96, 186–196. doi: 10.1038/labinvest.2015.142

Wong, M., Hendrix, M. J. C., Von Der Mark, K., Little, C., and Stern, R. (1984). Collagen in the egg shell membranes of the hen. Dev. Biol. 104, 28–36. doi: 10.1016/0012-1606(84)90033-2

Yang, W., Sherman, V. R., Gludovatz, B., Schaible, E., Stewart, P., Ritchie, R. O., et al. (2015). On the tear resistance of skin. Nat. Commun. 6, 6649. doi: 10.1038/ncomms7649

Yasser A. Ahmed and Soha A. Soliman. (2013). Long Bone Development in the Japanese Quail (Coturnix coturnix japonica) Embryos. Pak. J. Biol. Sci. 16, 911–919. doi: 10.3923/pjbs.2013.911.919

Yokomizo, T., Yamada-Inagawa, T., Yzaguirre, A. D., Chen, M. J., Speck, N. A., and Dzierzak, E. (2012). Whole-mount three-dimensional imaging of internally localized immunostained cells within mouse embryos. Nat. Protoc. 7, 421–431. doi: 10.1038/nprot.2011.441

Young, R. D., Knupp, C., Koudouna, E., Ralphs, J. R., Ma, Y., Lwigale, P. Y., et al. (2019). Cell-independent matrix configuration in early corneal development. Exp. Eye Res. 187, 107772. doi: 10.1016/j.exer.2019.107772

Young, R. D., Knupp, C., Pinali, C., Png, K. M. Y., Ralphs, J. R., Bushby, A. J., et al. (2014). Three-dimensional aspects of matrix assembly by cells in the developing cornea. Proc. Natl. Acad. Sci. U.S.A. 111, 687–692. doi: 10.1073/pnas.1313561110

